# Interpretable deep learning to uncover the molecular binding patterns determining TCR–epitope interactions

**DOI:** 10.1101/2022.05.02.490264

**Authors:** Ceder Dens, Wout Bittremieux, Fabio Affaticati, Kris Laukens, Pieter Meysman

## Abstract

The recognition of an epitope by a T-cell receptor (TCR) is crucial for eliminating pathogens and establishing immunological memory. Prediction of the binding of any TCR–epitope pair is still a challenging task, especially for novel epitopes, because the underlying patterns are largely unknown to domain experts and machine learning models. To achieve a deeper understanding of TCR–epitope interactions, we have used interpretable deep learning techniques to gain insights into the performance of TCR–epitope binding machine learning models. We demonstrate how interpretable AI techniques can be linked to the three-dimensional structure of molecules to offer novel insights into the factors that determine TCR affinity on a molecular level. Additionally, our results show the importance of using interpretability techniques to verify the predictions of machine learning models for challenging molecular biology problems where small hard-to-detect problems can accumulate to inaccurate results.

## Background

When a pathogen enters the human body, antigen-presenting cells display short peptides of the pathogen (called epitopes) on their cell surface using a major histocompatibility complex (MHC). A T-cell receptor (TCR) sequence on the surface of a T-cell recognizes the epitope to activate and subsequently to initiate the adaptive immune response. Quasi-random genetic rearrangements of the V, D, and J genes that express the TCR sequence make the recognition of a large variety of epitopes possible. The activation of the T-cell is crucial for eliminating the pathogen and creating the immunological memory to prevent severe symptoms with future infections of the same pathogen (1).

Predicting the binding of any given TCR–epitope pair can potentially lead to many advances in healthcare, by aiding diagnostics, vaccine development, and cancer therapies. Although some machine learning tools that perform TCR-epitope binding predictions exist, their performance is still insufficient for many applications. Because the CDR3 region of the TCR sequence is in close contact with the epitope (2,3), for simplicity this is often used by prediction tools instead of the full TCR sequence (4–14). However, it is still unclear which underlying patterns within the CDR3 and epitope sequences lead to binding. Indeed, high-quality machine learning models that are able to learn relevant rules underlying TCR–epitope binding do not currently exist. Broadly, a distinction can be made between the seen-epitope and the unseen-epitope prediction task. For seen-epitope prediction, one attempts to predict if a TCR will bind a known epitope, which means that the training dataset includes TCR–epitope pairs that involve this epitope. This requires a much lower amount of generalization capabilities and pattern learning from the model because it can compare the CDR3 sequence from the test sample to previously seen samples with the same epitope. In contrast, the unseen-epitope prediction task is more difficult because the model is evaluated on samples with novel epitopes that have not been seen during training. This no longer allows matching against known sequences and requires the model to learn more general binding patterns to make a correct prediction. As a consequence, the state-of-the-art performance on the unseen-epitope task is much lower but patterns learned by those models are more interesting. They also have a higher disruptive potential for healthcare as the possible epitope space far exceeds any feasible database. However, due to the increased complexity of the prediction problem, methods that have been developed to tackle the unseen-epitope task have turned out to be complex machine learning models. All current methods are exclusively based on neural networks, which are notoriously poor at explaining why a prediction is made the way it is, and thus what recognition patterns may underlie an interaction.

Feature attribution extraction methods provide information on which input features are mainly used by machine learning models to make a prediction for a single sample (15). When extracting feature attributions from neural networks, we can divide the extraction methods in two classes. The first class consists of gradient-based methods. Neural networks typically consist of multiple hidden layers connecting the input layer with the output layer. The output of each inner layer becomes the input of its subsequent layer, passing the input through all layers. Each of these layers have weights that are learned when the model is trained. The weights define the relation between the input and output of a layer. The partial derivative with respect to one of the input features of all these weights combined is the gradient for that feature. Intuitively, one might think about the gradient of a feature as the amount of change in the predicted output for a minor change in the input feature. Vanilla (16) is the oldest and simplest gradient-based method. It gives each input feature an attribution by calculating the gradients of the neural network weights for the given input sample. Another method that is often used is Integrated Gradients (IG) (18). IG is a path-attribution method: it integrates the Vanilla gradients over a range of input samples. The input samples are constructed by taking samples on the linear interpolation path between a baseline input (e.g., 0 for all features) and the original sample. A third gradient-based method, SmoothGrad (SG) (19), is designed to reduce noise in the feature attributions by adding noise to the input. SG samples similar inputs by adding random noise to the original input and averages the Vanilla feature attributions for those samples. The second class of feature attribution extraction methods are model-agnostic methods. These treat the model as a black box and only require the ability to make predictions with a chosen input. One example is SHAP (20), which computes feature attributions by looking at the change in the predicted output probability when part of the feature values is replaced by their baseline value. In our case, the baseline is a distribution of actual input samples: during each iteration, a random feature value is chosen from the input distribution as the baseline. This method is repeated multiple times to average the influence of the randomly chosen baseline.

In this study, we have used interpretable deep learning techniques to understand TCR–epitope binding by applying feature attribution extraction methods to two state-of-the-art TCR–epitope prediction models: ImRex (4) and TITAN (5). ImRex is a convolutional neural network (CNN) that follows the general design of CNNs for image processing. ImRex converts CDR3 and epitope sequences into interaction maps by calculating the pairwise difference between selected physicochemical properties (hydrophobicity, hydrophilicity, mass, and isoelectric point) of the amino acids of both sequences. This interaction map can be interpreted as a multi-channel image, with each channel corresponding to a specific physicochemical property (Figure S1), after which TCR–epitope binding prediction is performed using a multi-layer CNN. TITAN is based on one-dimensional CNNs using a contextual attention mechanism. The CDR3 and epitope sequences are encoded using a trainable embedding and separately fed into multiple one-dimensional convolutional layers, followed by a context attention layer that uses the epitopes as context for the TCR sequences and vice versa. The attention weights of both sequences are concatenated, and a stack of dense layers is used to output the binding probability. Here, we have applied several feature attribution extraction methods to ImRex and TITAN to obtain insights into the performance of TCR–epitope binding prediction by state-of-the-art machine learning models and investigate the biological patterns that underlie TCR–epitope binding.

## Results

### Molecular distances underlie recognition patterns in TCR–epitope complexes

While the complete TCR sequence might influence the recognition of a given epitope, previous studies have shown that the CDR3 is the main binding region (2,3). This is illustrated in Figure 1 showing a bound TCR–epitope complex, with the CDR3 region a sequence of 14 amino acids and the epitope a sequence of 9 amino acids. A molecular interaction between two protein sequences can only occur when there is close contact between amino acids of both sequences. The distance between residues from both sequences can therefore be used as a measure for indicating which amino acids are likely responsible for the interaction. Furthermore, it can be expected that any model that attempts to predict this interaction must make use of these residues, and thus the pairwise distances between amino acids of the TCR and the epitope can be used as a metric for evaluating the learned patterns within a model. To this end, we collected 105 solved TCR–epitope structures from the public RCSB Protein Data Bank (PDB) database (22) as ground truth data.

**Figure 1.**
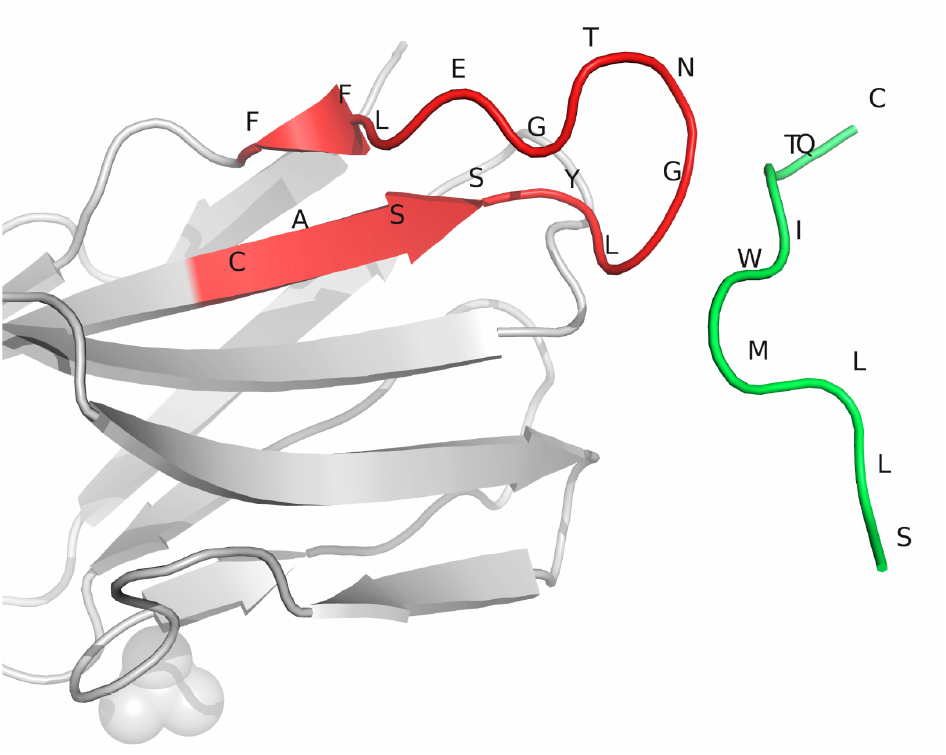
Molecular complex of a TCR–epitope interaction. The 3D image of the PDB complex 2P5W (21,22). Only the TCR beta chain and the epitope are shown, with the CDR3 region of the TCR beta chain colored red and the epitope colored green. The amino acids of both sequences are labeled with their one letter abbreviation.

### Feature attribution extraction methods reveal interacting residues in the ImRex model

We retrained the ImRex and TITAN models and evaluated them with epitope-grouped cross-validation, i.e. the train and test datasets do not share samples with the same epitope sequence. The performance is always given in the format: *metric*_*model*_ = *mean* ± *standard deviation*. Both models achieve an average receiver-operating characteristic (ROC) area under the curve (AUC) and precision-recall (PR) AUC of less than 56% (*ROC AUC*_*ImRex*_ = 0.550 ± 0.027; *ROC AUC*_*TITAN*_ = 0.559 ± 0.050; *PR AUC*_*ImRex*_ = 0. 556 ± 0. 033; *PR AUC*_*TITAN*_ = 0. 541 ± 0. 049), which shows that TCR–epitope binding prediction is still a very difficult task, even for state-of-the-art machine learning models. Given the low performance of these methods, it is unclear whether these models are truly learning the interaction. While it can be expected that these models may perform better with an increase in training data, a thorough evaluation of their current capacities is warranted. If these models are capturing the relevant molecular patterns, then they should be making use of those residues within the TCR and epitope sequences that are driving the interaction. Comparing the amino acid usage of the prediction tools with the actual distance between the amino acids may help to evaluate and to improve the performance and robustness of these tools.

To explore whether feature attribution extraction methods can be applied to these models, and which one is the most relevant, four common attribution extraction methods were applied to ImRex and TITAN. Extracting feature attributions for a sample from the ImRex model results in a 4-channel 2D feature attribution matrix with the same dimensions as the input sample. The attributions of the four physicochemical properties are summed per amino acid pair. For each pairwise combination of amino acids from the CDR3 and epitope sequence, the feature attribution extraction method returns a value that represents how much that input feature contributed to the prediction for the given input sample. For comparison to TITAN, the pairwise feature attributions from ImRex were merged per amino acid by taking the maximum of all feature attribution values for that amino acid. TITAN only uses the amino acid sequences as input, which results in one feature attribution value per amino acid. For each sample, we computed the Pearson correlation coefficient between the feature attributions and the residue proximity of the amino acids. The residue proximity is the inverted distance between the amino acids, as we expect an amino acid pair with a small distance to have a higher feature attribution. Also the pairwise residue proximity was merged per amino acid by taking the maximum value for each residue.

Figure 2 shows the correlation of each sample extracted with 4 different feature attribution extraction methods: Vanilla, Integrated Gradients (IG), SmoothGrad (SG), and SHAP from ImRex (fig. 2a and 2b) and TITAN (fig. 2c). For ImRex, there is a lower correlation when comparing pairwise feature attributions and residue proximity (fig. 2a) compared to per residue attributions and proximity (fig. 2b) for all four methods but the variation increases as well. Although less significant in the per residue case, feature attributions extracted with SG are most correlated with the residue proximity (*Corr*_*ImRex pairwise SG*_ = 0. 435 ± 0. 0175; *Corr*_*ImRexper residue SG*_ = 0. 516 ± 0.186). This can be explained by the de-noising effect of SG, as it will give higher attributions to the most important features and less to the other features. An additional explanation is that SG extracts the feature attributions multiple times with a slightly different input, which can make the algorithm more robust against high small-scale fluctuations in the gradient. For ImRex, we find that IG performs worse than Vanilla (*Corr*_*ImRexpairwise IG*_ = 0. 229 ± 0.117; *Corr*_*ImRexper residue SG*_ = 0. 384 ± 0. 230; *Corr*_*ImRex pairwise Vanilla*_ = 0. 318 ± 0.146; *Corr*_*ImRexper residue Vanilla*_ = 0. 474 ± 0.205). Feature attributions extracted by SHAP from ImRex are very weakly correlated with the residue proximity (*Corr*_*ImRex pairwise SHAP*_ = 0. 096 ± 0.102; *Corr*_*ImRexper residue SHAP*_ = 0. 214 ± 0.205), our results show that gradient based methods perform better. The feature attribution extraction methods on TITAN all perform similar and have only a limited improvement over the random correlation (*Corr*_*TITAN Vanilla*_ = 0. 299 ± 0.148; *Corr*_*TITAN IG*_ = 0. 297 ± 0.154; *Corr*_*TITAN SG*_ = 0. 342 ± 0.128; *Corr*_*TITAN SHAP*_ = 0. 293 ± 0. 116; *Corr*_*random*_ = − 0. 001 ± 0. 212). An overview of the correlation of all 9 feature attribution extraction methods we tested can be seen in Figure S2, a detailed ImRex feature attribution matrix for a single sample can be seen in Figure S3 and the feature attributions from different methods for TITAN can be seen in Figure S4.

**Figure 2.**
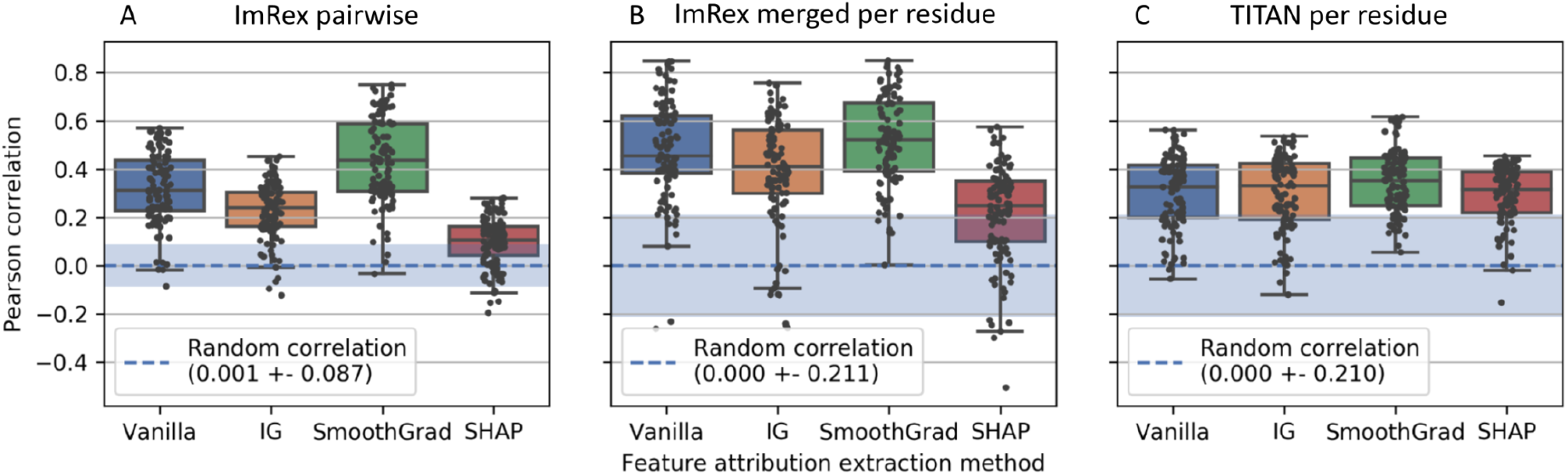
Pearson correlation coefficient of feature attributions and residue proximity extracted with different methods from ImRex and TITAN. The Pearson correlation is calculated between the feature attributions extracted with a specific method and the residue proximity between the amino acids of the TCR and epitope sequences. A boxplot is shown for each method giving the correlation over all 105 complexes. The random correlation is calculated by taking the Pearson correlation between a random feature attribution matrix and the actual residue proximity for each sample, repeated multiple times. Thus, this represents the correlation when the feature attribution extraction method would give a random output. Boxplots are constructed as follows: the box extends from the lower to upper quartile values of the data, with a line at the median. The whiskers extend from the box to the last datum before 1.5 times the interquartile range above/below the box. The random correlation is given as mean and standard deviation. **(A)** Correlation for ImRex using pairwise feature attributions and residue proximity. **(B)** Correlation for ImRex using feature attributions and residue proximity merged per residue. **(C)** Correlation for TITAN using feature attributions and residue proximity per residue.

### Feature attributions reveal important residues for each prediction

The remainder of all experiments were run using the SG feature attribution extraction method because it is the best scoring approach on both ImRex and TITAN.

The pairwise feature attributions from ImRex were merged per amino acid. For both ImRex and TITAN, all feature attributions were normalized per input sample by dividing them by the highest feature attribution value for that sample. This results in feature attributions ranging between 0 and 1, where the amino acid with the highest attribution gets a normalized value of 1, although the amino acid with the lowest attribution will not necessarily get a value of 0. The pairwise residue proximity is calculated for all pairs of amino acids from both sequences. A single value for each amino acid is derived in a similar way as for the pairwise feature attributions from ImRex: for each amino acid, the minimal distance (maximal proximity) to the amino acids from the other sequence is taken. The residue proximity is derived from the residue distance by taking 1/distance. At the end, the distance array is normalized per sample in the same way as the feature attributions. Figure 3 shows the feature attributions extracted from ImRex and TITAN with SG for the PDB complex 2P5W (21). For the epitope, we can see that the amino acids that are most important for TITAN are also in close contact with the CDR3 region, which is not the case for all important amino acids according to ImRex. ImRex focuses more on the residues at the beginning of the sequence. This can also be explained by the binding with the MHC molecule (see Discussion). The attributions for the CDR3 region are very different, where ImRex mainly focuses on the middle part of the CDR3 sequence while for TITAN all attributions are almost zero.

**Figure 3.**
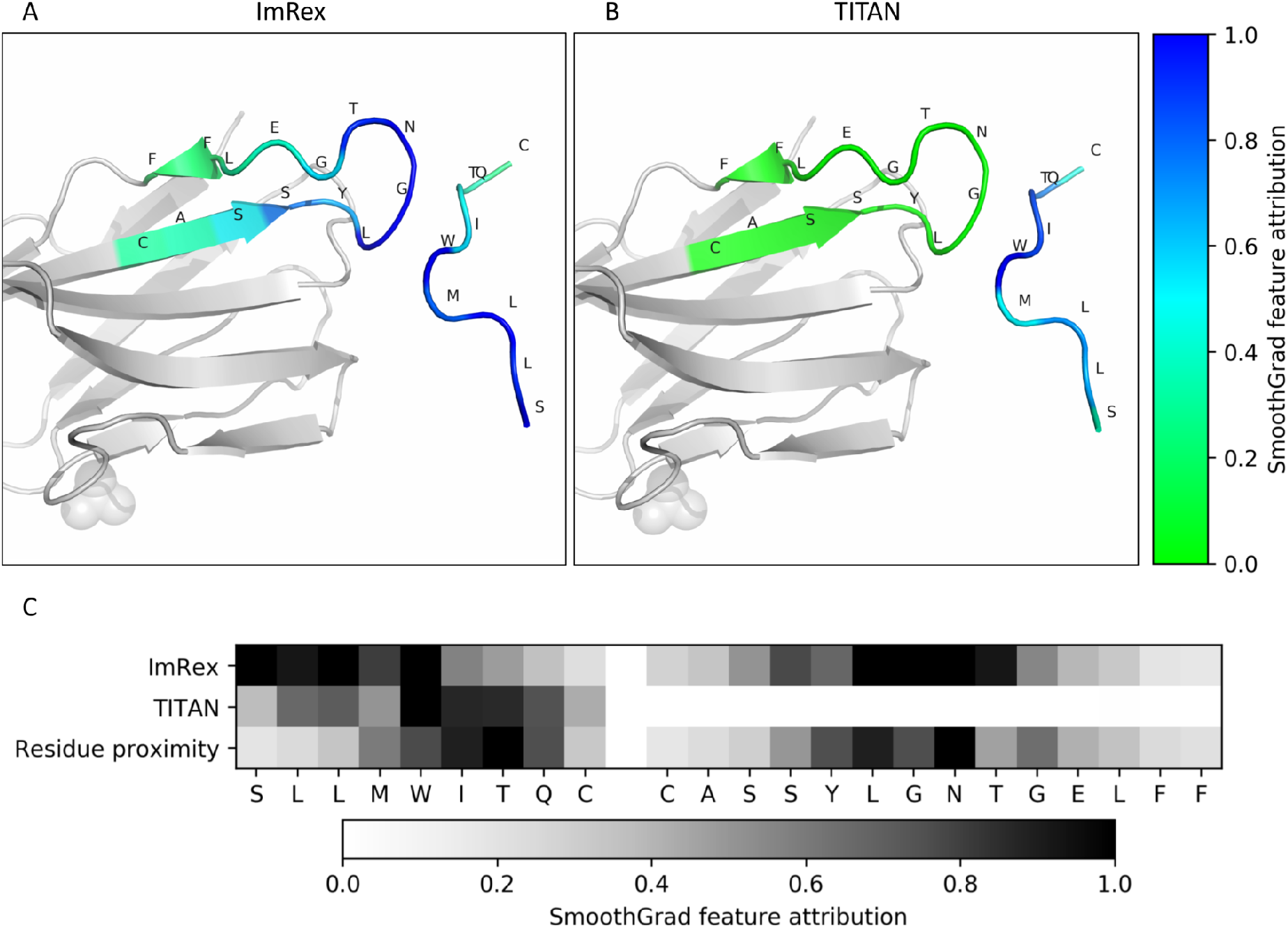
Feature attributions extracted with SG from PDB complex 2P5W. Feature attributions of **(A)** ImRex and **(B)** TITAN extracted with SG from the prediction for the PDB complex 2P5W (21,22) and shown on its molecular complex. Only the TCR beta chain and the epitope are shown. The TCR beta chain is colored gray, except for the CDR3 region which is colored according to a color range derived from the normalized feature attributions. The epitope is colored in the same way. **(C)** Feature attributions per position and model for the same complex together with the residue proximity. A higher value represents a larger feature attribution or residue proximity.

### Important residues are distinct for each TCR-epitope model

The findings derived from a single complex are representative for the full dataset, as can be seen in figure 4. We calculated the average attributions of the 105 samples and the average residue proximity. This shows that the amino acids of the epitope on position 6 to 9 (e_5_ - e_8_) are on average the closest to the CDR3 region. ImRex focuses more on the first part of the epitope while TITAN uses mostly the middle part. For the CDR3, the attributions from ImRex are similar to the 3D distance but more focused on the middle region. The average attributions from TITAN are very close to zero for each position of the CDR3 sequence, which suggests that TITAN primarily uses the epitope sequence to make its predictions.

**Figure 4.**
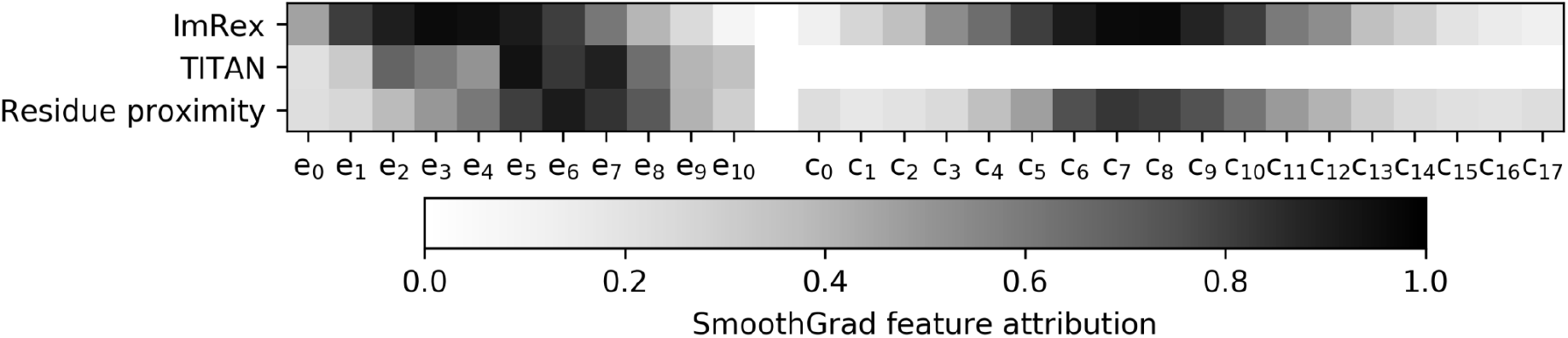
Average feature attributions per position. The average feature attribution per position and model extracted with SG, also the average residue proximity is given. Both the epitope and CDR3 sequence are padded left and right separately. The average is calculated by only looking at the feature attributions from sequences that do not have padding on that position.

We calculated the Pearson correlation coefficient between the feature attributions extracted from ImRex and TITAN and the proximity of the amino acids of the CDR3 and epitope sequence. The attributions of ImRex and the residue proximities were first merged per amino acid. This results in a higher correlation for ImRex when looking at both sequences together (*Corr*_*ImRex*_ = 0. 516 ± 0.186; *Corr*_*TITAN*_ = 0. 342 ± 0.128) (Figure 5). When only considering the epitope, the feature attributions from ImRex are not more correlated with the residue proximity than random attributions. This is caused by ImRex mainly focusing on the first residues of the epitopes. TITAN mainly uses the residues close to the CDR3 sequence (*Corr*_*ImRex ep*_ = 0. 045 ± 0. 446; *Corr*_*TITAN ep*_ = 0. 639 ± 0. 181). On the other hand, the correlation for the attributions of the CDR3 sequence is very high for ImRex and similar to random for TITAN (*Corr*_*ImRex CDR*3_ = 0. 807 ± 0. 104; *Corr*_*TITAN CDR*3_ = − 0. 187 ± 0. 301). This can be explained by the fact that TITAN always gives a near-zero attribution to the entire CDR3 sequence.

**Figure 5.**
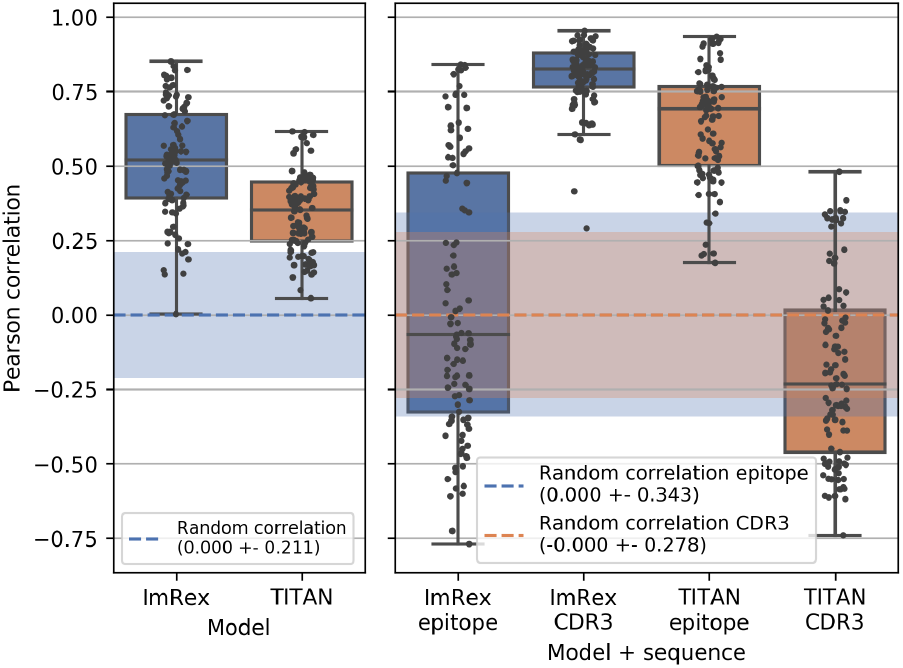
Pearson correlation between attributions and residue proximity for models and sequences. The correlation is calculated between the feature attributions extracted from ImRex and TITAN with SG and the proximity of the amino acid pairs from both sequences. For ImRex, the attributions were first merged per amino acid to allow comparison with TITAN. On the right, the correlation for both the epitope and CDR3 sequences separately are shown. The random correlation is calculated by taking the correlation between a random feature attribution array and the actual residue proximity for each sample, repeated multiple times. Thus, this represents the correlation when the feature attribution extraction method would give a random output. Boxplots are constructed as follows: the box extends from the lower to upper quartile values of the data, with a line at the median. The whiskers extend from the box to the last datum before 1.5 times the interquartile range above/below the box. The random correlation is given as mean and standard deviation.

### Feature attributions explain model performance

We re-evaluated both the ImRex and TITAN models on the task they were designed for: TCR–epitope interaction prediction. We use epitope-grouped cross-validation for all models which means that none of the epitopes will occur in samples of both the train and test split. Thus for these performance metrics, all test samples concern an unseen epitope. The default TITAN model has a slightly better average ROC AUC and ImRex has a slightly better average PR AUC (*ROCAUC*_*ImRex*_ = 0.550 ± 0.027; *ROCAUC*_*TITAN*_ = 0.559 ± 0. 05; *PR AUC*_*ImRex*_ = 0. 556 ± 0. 033; *PR AUC*_*TITAN*_ = 0. 541 ± 0. 049). The variance of both metrics is higher for TITAN, which also exhibited cross-validation splits with a performance below 0.500 (Figure 6). We previously found that TITAN primarily considers the epitope sequence to make its prediction. Here we see that it is still able to get a decent performance when evaluated with epitope-grouped cross-validation, even though a model that only uses the epitope sequence is not expected to achieve a good performance in this setting. We investigated this in more detail by training and evaluating the TITAN model on exactly the same data and cross-validation splits as ImRex (the ‘TITAN on ImRex data’ model in figure 6, also see figures S5 and S6 for average feature attributions and correlation of this model). When the TITAN model is trained on the same data as ImRex, the performance on both metrics drops (*ROC AUC*_*TITAN on ImRex data*_ = 0. 523 ± 0. 013; *PR AUC*_*TITAN on ImRex data*_ = 0. 514 ± 0. 012). This indicates that there might be an information leakage issue in the TITAN training data, which explains why a model that completely focuses on the epitope is still able to get a good performance on an unseen-epitope task. At last, we trained and evaluated the TITAN model on a third dataset with scrambled CDR3 sequences. This dataset is the same as the original dataset (including cross-validation train-test splits), but the CDR3 sequences are randomized by sampling random amino acids to create a new sequence of the same length for each original sequence, while ensuring that the distribution of the amino acids from the original CDR3 sequences is retained. The performance of the TITAN model trained on this random data is very similar to the performance of the original TITAN model (*ROC AUC*_*TITAN scrambled TCRs*_ = 0.559 ± 0. 045; *PR AUC*_*TITAN scrambled TCRs*_ = 0. 540 ± 0. 046), which is only possible when the CDR3 sequence does not contribute to the predictions. Thus, this confirms that our feature attribution extraction method made correct conclusions about the usage of the CDR3 sequence.

**Figure 6.**
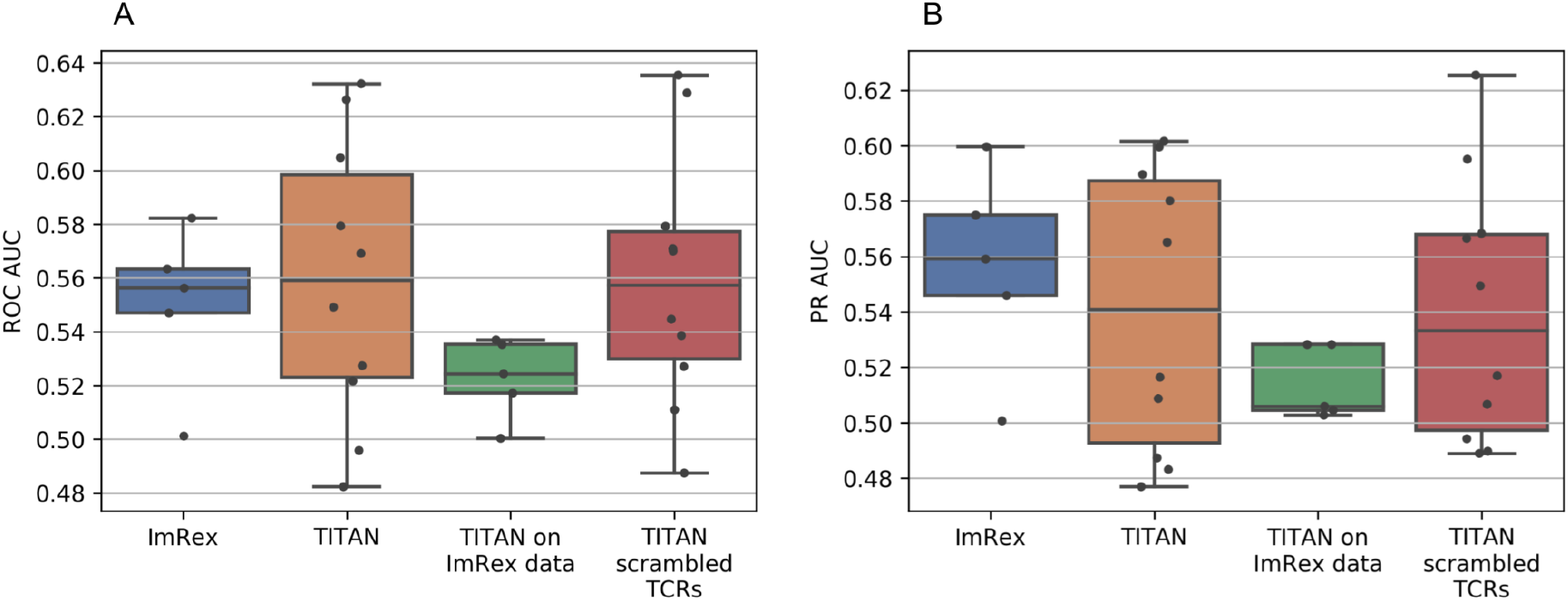
Comparison of model performance. The TCR–epitope interaction prediction performance of the different models measured with **(A)** ROC AUC and **(B)** PR AUC. ‘ImRex’ and ‘TITAN on ImRex data’ were both trained on the ImRex dataset and evaluated with 5-fold epitope-grouped cross-validation. ‘TITAN’ was trained on the original TITAN dataset and evaluated with 10-fold epitope-grouped cross-validation, ‘TITAN scrambled TCRs’ was trained on the TITAN dataset with scrambled CDR3 sequences and also evaluated with 10-fold epitope-grouped cross-validation. Boxplots are constructed as follows: the box extends from the lower to upper quartile values of the data, with a line at the median. The whiskers extend from the box to the last datum before 1.5 times the interquartile range above/below the box.

## Discussion

Although recently multiple improvements have been made in unseen-epitope TCR interaction prediction, the performance of the current state-of-the-art models is still limited (4,5,12). One of the reasons is that the determining factors for these molecular interactions are still unknown to both human experts and machine learning models. We presented a method that can extract which amino acids of the input were mainly used by the models to make their prediction. Highlighting those feature attributions on the molecular complex gives additional information to the domain expert about why the prediction was made and can give new insights in the factors that determine TCR affinity on a molecular level.

Using the actual distance between the amino acids of the TCR and epitope sequences as ground truth, we were able to compare the performance of different feature attribution extraction methods and prediction models. We found that SG is the best method for ImRex and TITAN. This is not unexpected because it is designed to reduce noise, making it focus more on the most important residue pairs and less on the others. This is similar to the residue proximity; a limited number of residues from both sequences are very close to each other, the proximity between the other residues decreases quickly. Another reason can be that SG samples multiple similar inputs with random noise, this should make the algorithm more robust against high small-scale fluctuations in the gradient. On ImRex, all gradient-based methods perform significantly better than SHAP. We reason that these methods are better at finding more detailed attributions because they have access to the gradients of the models while SHAP only compares the predicted output for altered inputs.

We found that, on average, ImRex uses mainly the start and middle amino acids of the epitope, while the distance is smaller for amino acids on position 6 to 9 (out of 11). This points to a first limitation of our study, as we only considered the distance between the TCR and epitope. However, one must also consider the interaction between the epitope and the MHC molecule, which is known to occur close to either end of the epitope for MHC class I. ImRex is revealed to use the amino acids of the CDR3 region similarly to the average residue proximity with somewhat more attribution to the middle part and less attribution to the outer parts of the sequence. For TITAN, we found that the amino acid usage of the epitope sequence is similar to the residue proximity, but it does not consider the CDR3 sequence. However, when testing TITAN using epitope-grouped cross-validation, unexpectedly its performance was still similar to ImRex. Training TITAN on the ImRex data resulted in a much lower performance, which leads us to hypothesize that the unexpected performance can be explained by how the TITAN training and evaluation data was constructed. TITAN creates its negative samples in a different way than ImRex. TITAN uniformly samples a random epitope for each TCR sequence, while ImRex samples a random epitope with the same probability as in the positive dataset. This can be seen in supplementary table S1: for each epitope the number of positive samples is between 15 and 400 (which is expected due to the data preprocessing), but the number of negative samples is similar for all epitopes. This leads to a large imbalance between the number of positive and negative samples for most epitopes. Note that the same imbalance is also present in the training data (although with other epitopes). This imbalance is learned by the TITAN model that thereby completely focuses on the epitope sequence, as we found by extracting the feature attributions. The last column of supplementary table S1 shows that TITAN almost always gives a negative prediction for most epitopes except for a few (TLIGDCATV, RQLLFVVEV, EPLPQGQLTAY, and LSDDAVVCFNSTY in this specific cross-validation split) for which it almost always gives a positive prediction.

Even though the dataset is imbalanced, this does not fully explain the good test performance. TITAN splits its data in two datasets, a train and test set, where no epitopes are shared among each. The model is trained on the train dataset and at the end of each epoch tested on the test dataset. When retraining the model ourselves, we saw that the performance on the test dataset is very unstable across multiple epochs and does not converge. After training for a given number of epochs, the test performance of the best epoch is selected and reported as the final test performance. We therefore hypothesize that the model tries to give a high prediction to random epitopes or patterns in the train data, repeatedly changing this every epoch (which results in the very high variation in performance across epochs). At the end, the epoch where the patterns selected from the train dataset gave (by chance) the best performance on the test dataset is chosen. If true, this could have been avoided by using an independent validation dataset to determine the best epoch. Afterwards, the performance could have been given by applying the model from that epoch on the test set.

## Conclusions

We have shown that applying feature attribution extraction methods are useful to improve the explainability for protein interaction prediction, as applied here to the TCR–epitope problem. The gradient based methods performed clearly better than the model-agnostic method SHAP. Showing feature attributions on the TCR–epitope molecular complex can be useful to gain more knowledge about why specific protein sequences interact. Extracting feature attributions is a good way to verify a model and data and to check that it works as expected. This is especially true for challenging problems, where small hard-to-detect issues in the dataset balance or evaluation methods can compound to inaccurate results.

## Methods

### Data

#### Molecular complex data

A collection of TCR–epitope MHC class I complexes with links to their RCSB Protein Data Bank (PDB) (22) entry was downloaded from the TCR3d database (23) and all non-human entries were removed. The TCR beta chain, epitope chain and location of the CDR3 region were manually selected from the PDB complexes with the help of the IMGT/3Dstructure-DB (24–26) online tool. Finally, we ended up with 105 unique complexes.

#### ImRex training data

We used the data that was also used in the ImRex (4) paper. It uses the VDJdb dataset from August 2019 (27) and is filtered on samples with human TCR beta sequences. All samples from the 10x Genomics study (28) were excluded and only samples with a length between 10-20 and 8-11 for respectively the CDR3 and epitope were kept. The data was downsampled to have at most 400 samples per epitope and negative data was generated by shuffling. We performed one additional filtering step: all samples that are also present in the molecular complex data (based on the CDR3 sequence) were removed, which reduced the positive dataset size from 6702 to 6656 samples. Each model trained on the ImRex data was evaluated with 5-fold epitope-grouped cross-validation, as per the original study. This divides the data in five groups with about the same amount of epitopes and a similar distribution of number of samples per epitope. Samples with the same epitope are all put in the same group.

#### TITAN training data

For TITAN (5), we use their ‘strictsplit’ data which is a combination of two datasets: the VDJdb (27) and a COVID-19 specific dataset published by the ImmuneCODE project (29). Both datasets were filtered on human TCR beta sequences, all epitopes with less than 15 associated TCRs were removed and the data was downsampled to have at most 400 samples per epitope. For the COVID-19 dataset, only samples with a single unique epitope were kept and unproductive samples were excluded. After merging both datasets, negative samples were generated by shuffling. This means that a negative sample is generated for each positive sample, pairing the original CDR3 sequence with a random, different epitope from the positive dataset. The only additional filtering step we performed on the data was removing all samples that are also present in the molecular complex data, which reduced the positive dataset size from 23,145 to 23,125 samples.

We also created an additional dataset with scrambled CDR3 sequences. We used the final TITAN dataset but replaced all CDR3 sequences with a random combination of amino acids of equal length. The amino acid distribution of the CDR3 sequences of the original dataset was kept.

Models trained on any of the TITAN datasets are always evaluated with 10-fold epitope-grouped cross-validation. The 10 groups created by TITAN were kept.

### ImRex model training

We retrained ImRex with the same parameters as the final published model (30). This is the default ‘padded model’, has a batch size of 32, is trained for 20 epochs, uses ReLU activation functions, has convolutional layers with depth 128, 64, 128, and 64, has a dropout rate of 0.25 for the convolutional layers, uses a learning rate of 1e-4, a regularization of 0.01 on all layers and uses the RMSProp optimizer (31).

### TITAN model training

TITAN was trained with the parameter configuration of the AA CDR3 case as explained in their paper (the configuration on their online repository led to similar results). We set the padding of the CDR3 and epitope to 25 instead of 500, otherwise we could not achieve a better than random performance. The epitope is not encoded as SMILES (32), the size of the dense hidden layers is 368 and 184, a ReLU activation function is used, a dropout of 0.5, a batch size of 512 without normalization, a learning rate of 1e-4, the attention size for both sequences is 16, the embedding size for both sequences is 26, the epitope and CDR3 kernel sizes are both [3, 26], [5, 26], and [11, 26] and the epitope and CDR3 embeddings are both learned during training.

Retraining TITAN on its own data was done with the original 10-fold epitope-grouped cross-validation splits provided by the authors. Retraining TITAN on the ImRex data was done with the 5-fold epitope-grouped cross-validation splits generated by ImRex.

### Feature attribution extraction

The input for ImRex can be represented as an image, so it is possible to apply feature attribution extraction methods made for image classification CNNs. These methods give an attribution to each of the input pixels for the prediction of a single sample. In our case, every pixel is a pairwise combination of an amino acid from both sequences. The value of the attribution represents how much each pixel contributed to the prediction.

In this research, we focused on two kinds of feature attribution extraction methods: a set of methods based on the neural network gradients and a model-agnostic method: SHAP (20). Gradient based methods can only be used on models with a differentiable input, whereas SHAP can be used for any type of black box model. We compared 8 different feature attribution extraction methods that are based on gradients (Vanilla (16), Integrated Gradients (IG) (18), SmoothGrad (19), SmoothGradIG (18,19), GuidedIG (18,33), BlurIG (18,34), SmoothGradBlurIG (18,19,34) and XRAI (35)) and the SHAP method using random sampling from the dataset as background. Of these, for conciseness, four representative methods were used in the results section (Vanilla, IG, SmoothGrad and SHAP), while results for all methods are available as supplementary information. The Python package *shap* (version 0.40.0) (20) was used for the implementation of the SHAP algorithm and the other methods were implemented with the Python package *saliency* (version 0.1.3) (16,19,33–36).

TITAN uses a 1D categorical input: a concatenated list of the amino acids from the epitope and CDR3. Therefore, there will only be one feature attribution value for each amino acid. Because gradient based feature attribution extraction methods require a differentiable input, the embedded input was used with TITAN.

### Feature attribution evaluation

We evaluated the feature attribution extraction methods by comparing them to the distance between the amino acids of both sequences in the molecular complex. The distance between two amino acids is the minimal distance between two atoms of each amino acid in ångströms. This results in a distance matrix with values between 2.3Å and 30Å. The amino acids that are closer to the other sequence are also more important for the interaction. Therefore, we inverted each value of the distance matrix by taking 1/value to get the residue proximity. The feature attributions and the residue proximity matrix are both normalized by dividing them by their maximum value. This results in all values being between 0 and 1 and the largest value is always equal to 1.

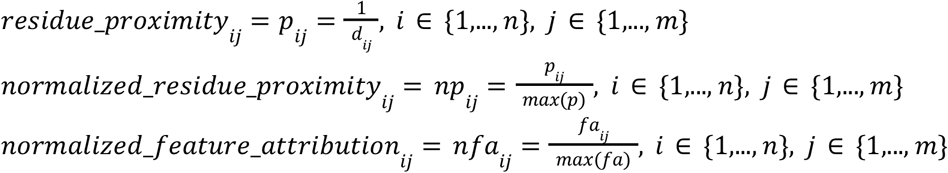

With n the length of the epitope, m the length of the CDR3 sequence, *d*_*ij*_ the distance between amino acid i of the epitope and amino acid j of the CDR3 sequence and *fa*_*ij*_ the feature attribution of the amino acid pair (i, j).

To quantify the correlation between the feature attributions and the residue proximity the Pearson correlation coefficient was calculated between the feature attributions and the residue distance for each sample and multiplied by -1. This results in a higher, positive value when there is a higher correlation between the feature attributions and the residue proximity.

### 1D feature attributions

To compare the feature attributions and the correlation with the residue proximity between ImRex and TITAN we converted the 2D ImRex attributions and distance matrices to a 1D array by merging the values per amino acid and concatenating the result of both sequences. For each amino acid the maximum feature attribution and minimum distance with respect to every amino acid from the other sequence was taken. Those 1D arrays (of length *n* + *m*, with n the length of the epitope and m the length of the CDR3) were inverted and normalized in the same way as the 2D matrices.

### Molecular complex highlighting

The normalized 1D feature attributions from both models are shown on the molecular complex with PyMol (version 2.3.0) (37). For clarity, we only show the TCR beta sequence and the epitope. The TCR beta chain is colored gray, except the CDR3 region which is colored according to a color range derived from the normalized feature attributions, green for a feature attribution of zero to blue for a feature attribution of one. The epitope is colored with the same color range. The amino acids of both sequences are labeled with their one letter abbreviation.

## Declarations

### Data and Code Availability

The datasets analyzed during the current study and all scripts used to obtain the results are licensed under the Apache License 2.0 and are available on GitHub at https://github.com/PigeonMark/McFAE (38) and on Zenodo at https://doi.org/10.5281/zenodo.6500495 (39). All code is written in Python 3.7 (40). PyTorch (version 1.10.0) (41), TensorFlow (GPU version 2.6.0) (42) and Keras (version 2.6.0) (43) were used for implementing the neural network architectures and model training. NumPy (version 1.18.1) (44) and pandas (version 1.0.1) (45,46) were used for data processing. SHAP (version 0.40.0) (20) was used to extract SHAP feature attributions and Saliency (version 0.1.3) (36) was used to extract the other feature attributions. Matplotlib (version 3.1.3) (47), seaborn (version 0.10.0) (48), and Pillow (version 7.0.0) (49) were used to create the figures. PyMol (version 2.3.0) (38) was used to create the molecular complex figures.

### Authors’ contributions

CD performed the study and wrote the manuscript. FA performed a preliminary study on applying integrated gradients on ImRex. WB, KL and PM conceived and supervised the study and revised the manuscript. All authors read and approved the final manuscript.

### Materials & Correspondence

Correspondence to Pieter Meysman.

