## Supplementary figures and tables for "Interpretable deep learning to uncover the molecular binding patterns determining TCR–epitope interactions"

### Supplementary Material

#### Supplementary Figures

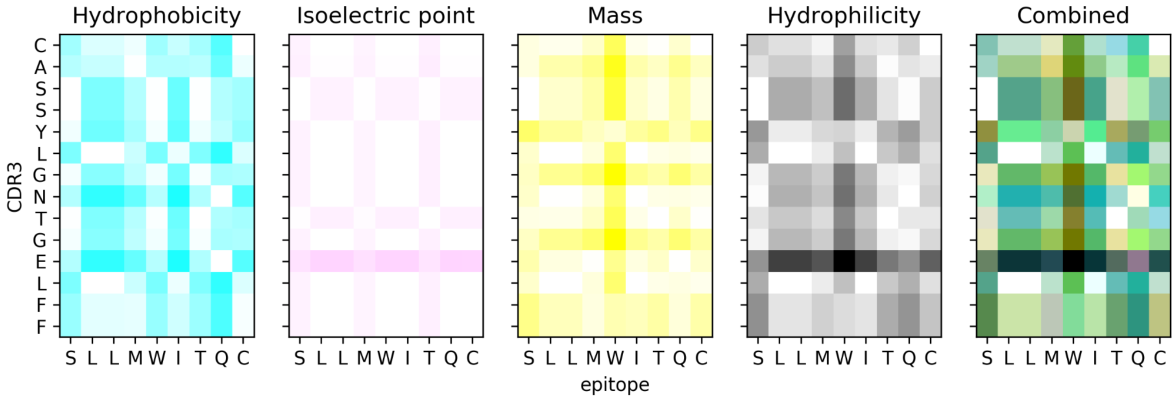


**Figure S1. ImRex feature encoding.** ImRex converts its input to an interaction map by calculating the absolute pairwise differences of four physicochemical properties: Hydrophobicity, Isoelectric point, Mass, and Hydrophilicity. Each of these features can be represented as a channel of a CMYK image. The feature values for PDB complex 2P5W are shown, the combined CMYK image is shown on the right.

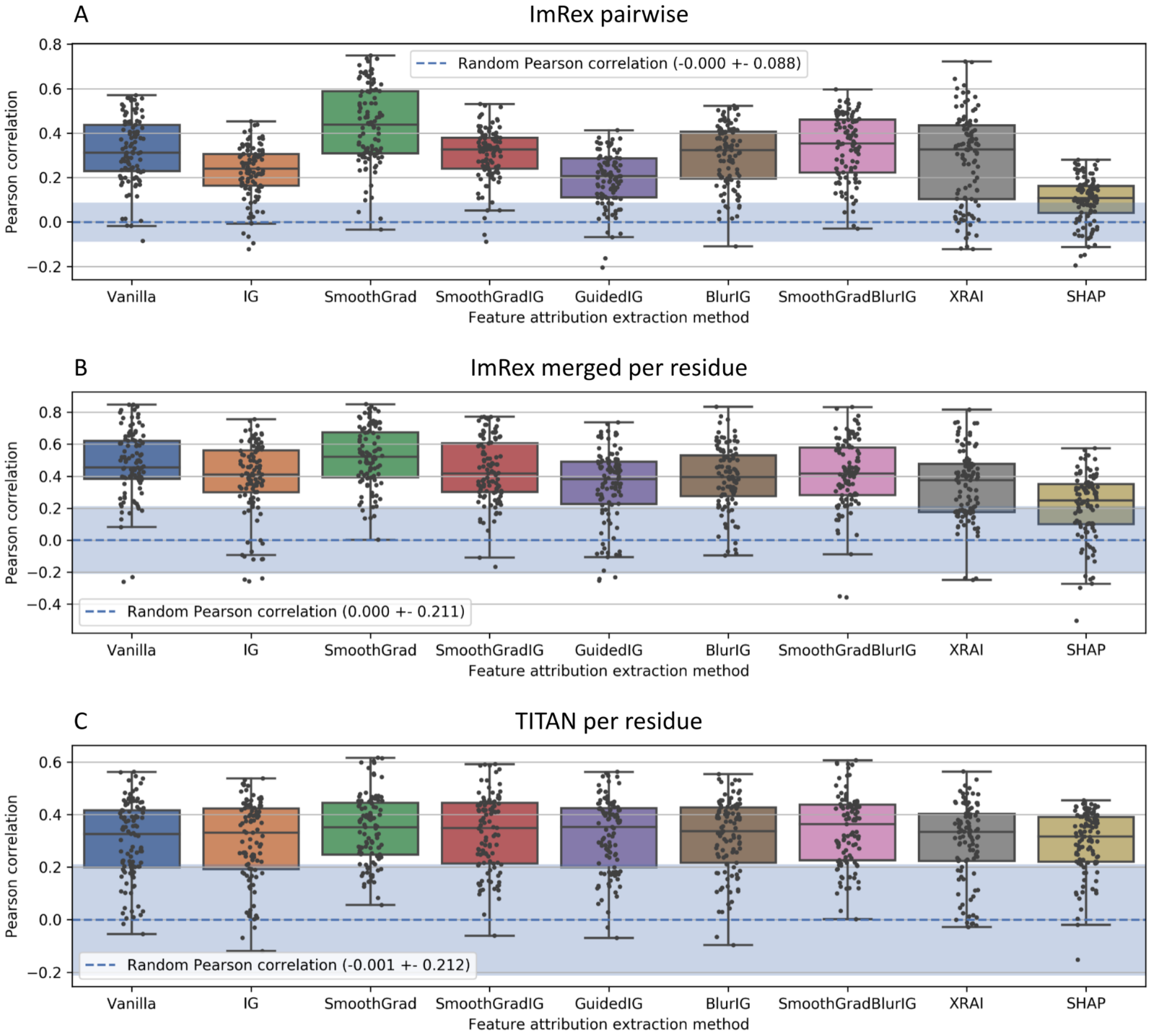


**Figure S2. Pearson correlation coefficient of feature attributions and residue proximity extracted with different methods from ImRex and TITAN.** The Pearson correlation is calculated between the feature attributions extracted with a specific method and the residue proximity between the amino acids of the TCR and epitope sequences. A boxplot is shown for each method giving the correlation over all 105 complexes. The random correlation is calculated by taking the Pearson correlation between a random feature attribution matrix and the actual residue proximity for each sample, repeated multiple times. Thus, this represents the correlation when the feature attribution extraction method would give a random output. Boxplots are constructed as follows: the box extends from the lower to upper quartile values of the data, with a line at the median. The whiskers extend from the box to the last datum before 1.5 times the interquartile range above/below the box. The random correlation is given as mean and standard deviation. **(A)** Correlation for ImRex using pairwise feature attributions and residue proximity, **(B)** Correlation for ImRex using feature attributions and residue proximity merged per residue, and **(C)** Correlation for TITAN using feature attributions and residue proximity per residue. Vanilla, IG, SmoothGrad and SHAP were selected to be shown in the results. SmoothGradIG, GuidedIG, BlurIG, SmoothGradBlurIG and XRAI are all variations on SG and/or IG.

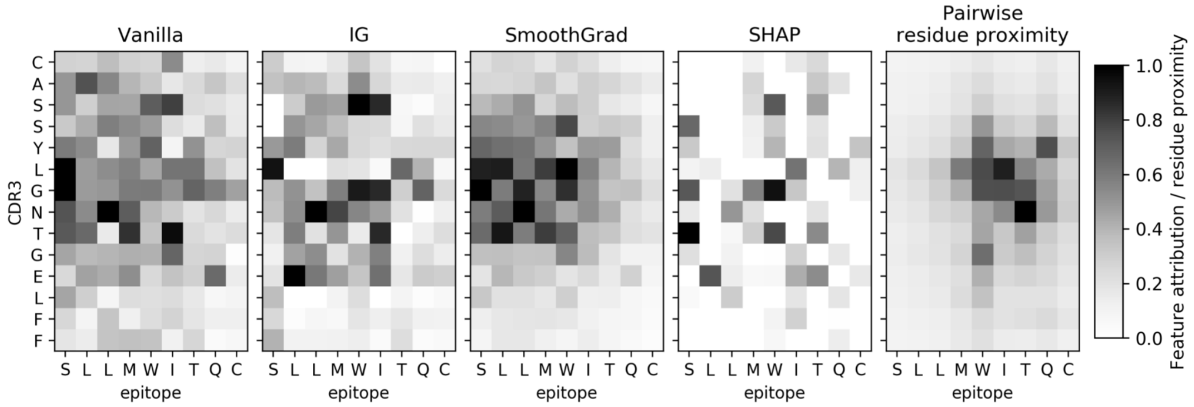


**Figure S3. Pairwise ImRex feature attributions.** Feature attributions extracted from ImRex for PDB complex 2P5W with different extraction methods: Vanilla, IG, SmoothGrad and SHAP. The feature attributions are first summed per amino acid pair to get a grayscale image. Amino acid pairs with a higher value (darker) received a higher feature attribution. On the right, the pairwise residue proximity is given, a higher value (darker) means that those amino acids are closer to each other.

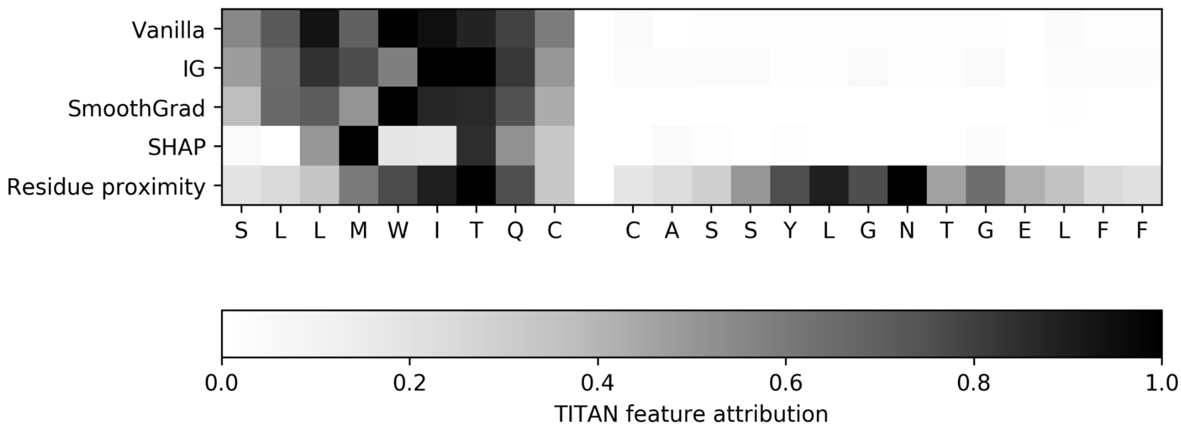


**Figure S4. TITAN feature attributions.** Feature attributions extracted from TITAN for PDB complex 2P5W with different extraction methods. The bottom row gives the residue proximity merged per amino acid. The CDR3 sequence gets an attribution of almost zero by all of the methods.

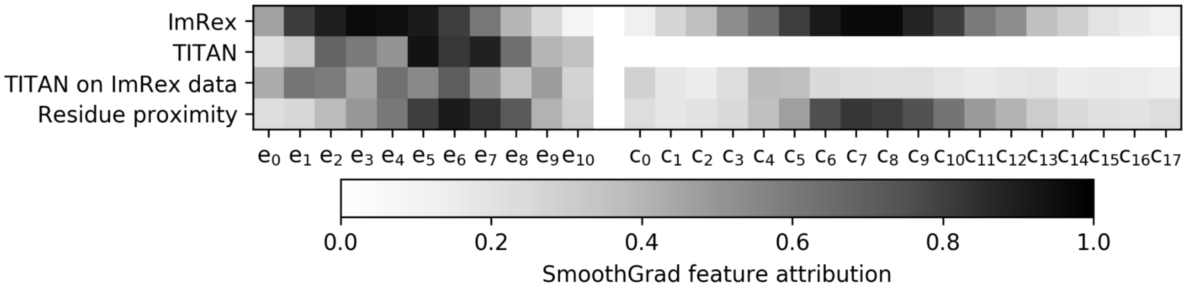


**Figure S5. Average feature attributions per position.** The average feature attribution per position and model, also the average residue proximity is given. Both the epitope and CDR3 sequence are padded left and right separately. The average is calculated by only looking at the feature attributions from sequences that do not have padding on that position. A higher value represents a higher feature attribution for ImRex, TITAN and TITAN on ImRex data or a higher residue proximity. TITAN trained on the ImRex data gives much less attribution to epitope and also slightly uses the first part of the CDR3 sequence for its predictions.

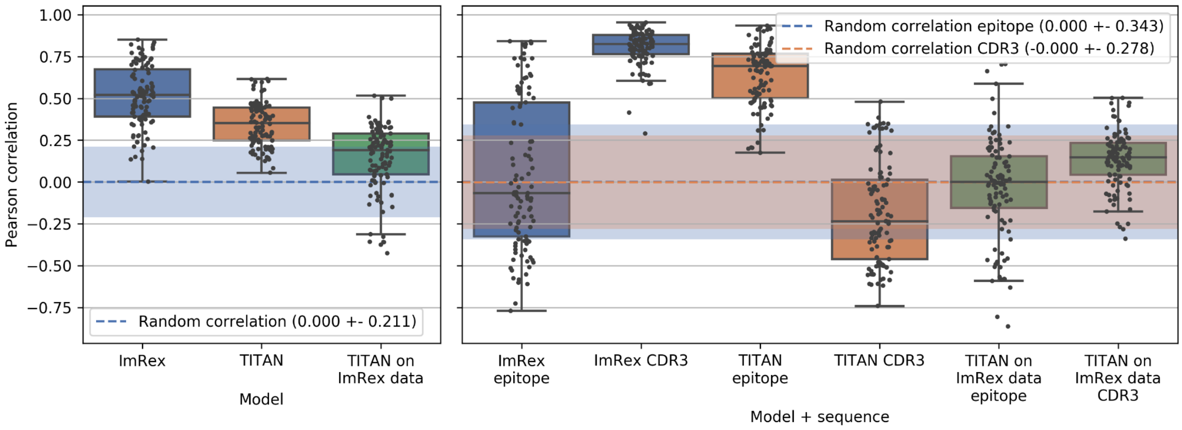


**Figure S6. Pearson correlation between attributions and residue proximity for models and sequences.** The correlation is calculated between the feature attributions extracted from ImRex, TITAN and TITAN on ImRex data with SG and the proximity of the amino acid pairs from both sequences. For ImRex, the attributions were first merged per amino acid to allow comparison with TITAN. On the right, the correlation for both the epitope and CDR3 sequences separately are shown. The random correlation is calculated by taking the correlation between a random feature attribution array and the actual residue proximity for each sample, repeated multiple times. Thus, this represents the correlation when the feature attribution extraction method would give a random output. Boxplots are constructed as follows: the box extends from the lower to upper quartile values of the data, with a line at the median. The whiskers extend from the box to the last datum before 1.5 times the interquartile range above/below the box. The random correlation is given as mean and standard deviation.

**Table S1. Example of TITAN dataset, statistics per epitope.**

| Epitope | Number of positive samples | Number of negative samples | Precision | Recall | Predicted positive ratio |
|---|---|---|---|---|---|
| TLIGDCATV | 399 | 109 | 0.786 | 1.000 | 1.000 |
| RQLLFVVEV | 399 | 117 | 0.780 | 0.853 | 0.845 |
| YLQPRTFLL | 304 | 93 | 0.000 | 0.000 | 0.000 |
| YLDAYNMMI | 236 | 107 | 0.000 | 0.000 | 0.000 |
| ALSKGVHFV | 181 | 115 | 0.000 | 0.000 | 0.000 |
| KTSVDCTMYI | 137 | 118 | 0.000 | 0.000 | 0.000 |
| IQYIDIGNY | 122 | 123 | 0.000 | 0.000 | 0.000 |
| IPSINVHHY | 92 | 130 | 0.000 | 0.000 | 0.000 |
| SLVKPSFYV | 83 | 139 | 0.000 | 0.000 | 0.022 |
| LVLSVNPYV | 62 | 111 | 0.000 | 0.000 | 0.000 |
| HPKVSSEVHI | 54 | 151 | 0.000 | 0.000 | 0.000 |
| ISDYDYYRY | 44 | 131 | 0.000 | 0.000 | 0.000 |
| GTHWFVTQR | 39 | 124 | 0.000 | 0.000 | 0.000 |
| EPLPQGQLTAY | 32 | 114 | 0.219 | 1.000 | 1.000 |
| EHPTFTSQYRIQGKL | 30 | 122 | 0.000 | 0.000 | 0.000 |

| | | | | | |
|---|---|---|---|---|---|
| FQPTNGVGY | 26 | 120 | 0.000 | 0.000 | 0.000 |
| CTELKLSDY | 24 | 125 | 0.000 | 0.000 | 0.000 |
| LSDDAVVCFNSTY | 18 | 139 | 0.107 | 0.889 | 0.937 |
| KMQRMLLEK | 18 | 112 | 0.000 | 0.000 | 0.000 |

Example overview of the TITAN unseen-epitope dataset, this is the test set of one cross-validation fold but test sets from all folds show similar trends. The number of positive/negative samples have a similar imbalance in all train datasets, the last three columns can not be computed for the train dataset. All samples from this dataset are sorted per epitope and the number of positive and negative samples per epitope are given. In the next columns, the precision and recall of the predictions are calculated using only the samples for that epitope. The last column gives the ratio of samples that were predicted to be positive for that epitope.
